# Impaired tumor death receptor signaling drives resistance to CAR T cell therapy

**DOI:** 10.1101/627562

**Authors:** Nathan Singh, Olga Shestova, Pranali Ravikumar, Katharina E. Hayer, Seok Jae Hong, Elena J. Orlando, Karen Thudium Mueller, Charly R. Good, Shelley L. Berger, Ophir Shalem, Matthew D. Weitzman, Stephan A. Grupp, Carl H. June, Saar Gill, Marco Ruella

**Author notes:** These authors contributed equally to this work (Saar Gill, Marco Ruella). Correspondence and requests for materials should be addressed to N.S., S.G., or M.R.

## Abstract

Cellular immunotherapy using T cells engineered to express chimeric antigen receptors targeting CD19 (CART19) leads to long-term remission in patients with B-cell malignancies^1–5^. Unfortunately, a significant fraction of patients demonstrate primary resistance to CART19 or experience relapse after achieving remission. Beyond loss of target antigen, the molecular pathways governing CART19 failure are unknown. Here we demonstrate that death receptors, cell surface signaling molecules that induce target cell apoptosis, are key mediators of leukemic resistance and CART19 failure. Using a functional CRISPR/Cas9-based genome-wide knockout screen^6^ in B-cell acute lymphoblastic leukemia (ALL), we identified the death receptor signaling pathway as a central regulator of sensitivity to CART19-induced cell death. In the absence of the pro-apoptotic death receptor signaling molecules BID or FADD, ALL cells were resistant to CART19 cytotoxicity, resulting in rapid disease progression in mice. We found that this initial resistance to cytotoxicity led to persistence of tumor cells, which drove the development of T cell dysfunction that further compromised anti-tumor immunity and permitted tumor outgrowth. We validated these findings using clinical samples collected from patients with ALL treated with CD19-targeted CAR T cells and found that expression of pro-apoptotic death receptor pathway genes in pre-treatment tumor samples correlated with CAR T cell expansion and persistence, as well as patient response and overall survival. Our findings indicate that tumor-intrinsic death receptor signaling directly contributes to CAR T cell failure.

## Main Text

The success of CD19-directed CAR T cells (CART19) is significantly limited by lack of disease response (primary resistance) and disease relapse after initial response (acquired resistance)^1–5^ in some patients. To identify the mechanisms that enable acquired resistance to CART19, we performed genome-wide CRISPR/Cas9-based knockout screens^6^ in the CD19+ human ALL cell line Nalm6 (**Figure 1a**). Using the Brunello short-guide RNA (sgRNA) library^7^, we generated a pool of Nalm6 in which each cell had been edited for loss of function of a single gene. We combined Brunello-edited Nalm6 cells with CART19 and monitored co-cultures for the emergence of resistant leukemic clones. GFP-labeled Nalm6 initially became undetectable after three days, indicating robust early cytotoxicity by CART19 cells, but recurred after 28 days (**Extended Data Figure 1**). Next-generation sequencing demonstrated that the single most-enriched guide target in resistant Nalm6 cells was *CD19* (**Figure 1b**). This finding is consistent with the clinical observation that antigen loss is the primary cause of acquired resistance (relapse) after initial CART19-induced remission^8–10^, and validated our screening approach.

**Figure 1.**
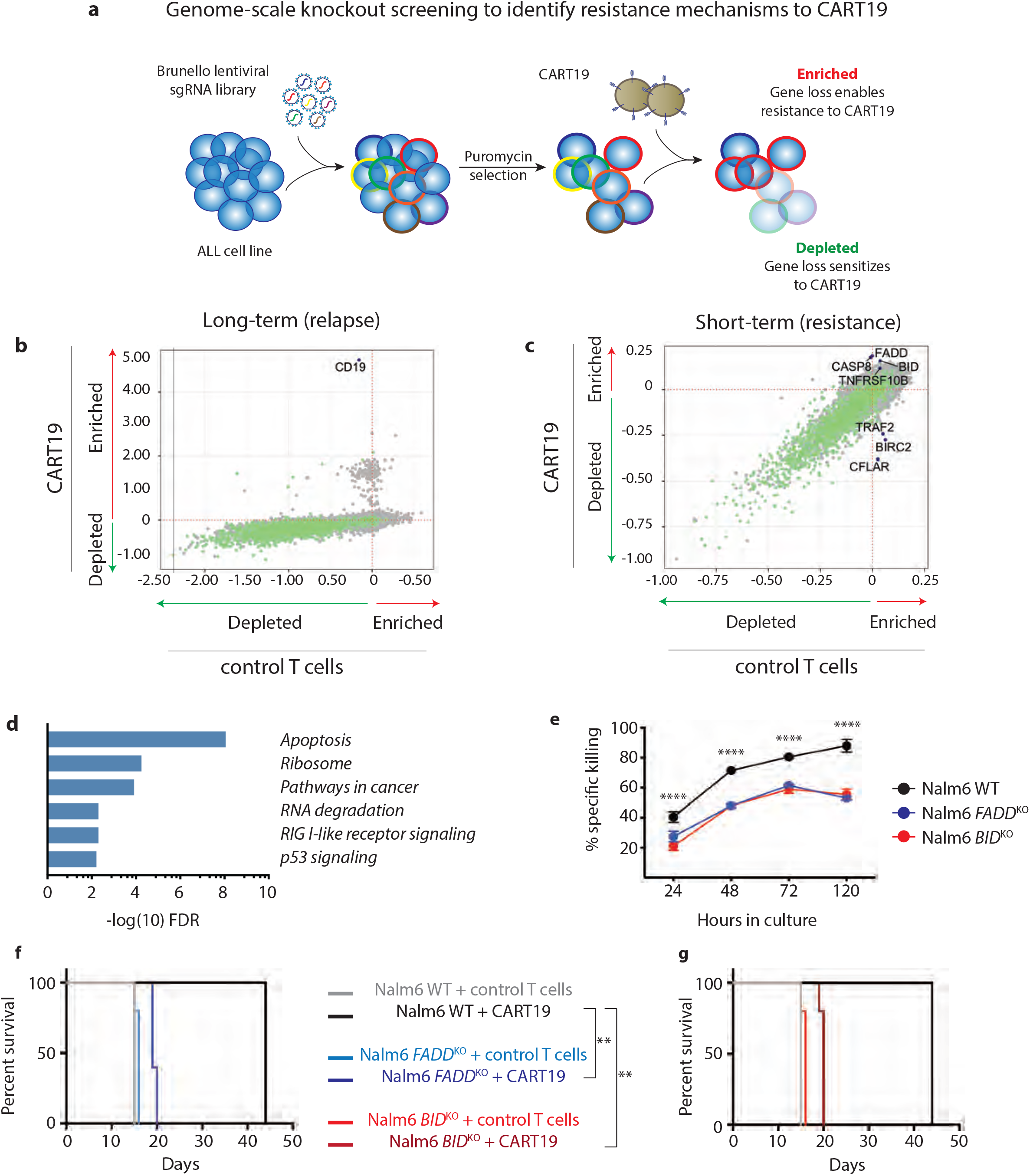
Death receptor signaling is a key mediator of resistance to CART19. **a,** Schematic of genome-wide knockout screen using the Nalm6 ALL cell line and CD19-targeted CAR T cells. **b-c,** Scatter plot of normalized MAGeCK algorithm beta scores representing enriched and depleted sgRNAs after **b,** long-term and **c,** short-term CART19 co-culture. Essential genes are identified in green^30^. **d,** Gene set enrichment analysis of the most-enriched and most-depleted sgRNAs identified in the short-term screen using the Kyoto Encyclopedia of Genes and Genomes (KEGG) pathway database. **e,** Nalm6 cells edited for knockout of either *FADD* or *BID* were exposed to CART19 cells. **f-g,** immunode-ficient mice were engrafted with WT, *FADD* or *BID* KO Nalm6 and given 5×10^5^ control T cells or CART19, and survival was measured over time. *P<0.05, **P<0.01, ***P<0.001 ****P<0.0001 by ANOVA (**e**) and Log-Rank test (**f-g**).

While several mechanisms of acquired resistance by antigen silencing have been described^9–11^, less is known about the mechanisms of primary resistance. Nearly 1 in 5 patients with CD19+ ALL are primarily refractory to CART19^1,3^, indicating that some tumors are intrinsically resistant to this therapy. To identify the processes responsible for primary resistance, we combined Brunello-edited Nalm6 with CART19 and collected cells after 24 hours of co-culture. In this short-term screening system, sgRNA sequencing demonstrated enrichment of sgRNAs targeting several pro-apoptotic death receptor signaling molecules (*FADD, BID, CASP8* and *TNFRSF10B*), and depletion of guides targeting several anti-apoptotic molecules (*CFLAR, TRAF2* and *BIRC2*) (**Figure 1c**). Gene set enrichment analysis (GSEA) identified a strong association between enriched or depleted sgRNAs and apoptotic signaling (false discovery rate *P*=1.13×10^-8^, **Figure 1d**). Notably, all sequenced sgRNAs in this pathway are specifically involved in death receptor-driven apoptosis. These findings suggest that ALL death receptor signaling is a key regulator of resistance to CART19.

To confirm the functional significance of impaired tumor death receptor signaling, we focused on two targets: Fas-associated protein with death domain (FADD), which serves as a membrane-proximal adaptor to all pro-apoptotic death receptors, and BH3 interacting-domain death agonist (BID), which is activated downstream of FADD and initiates mitochondrial activity that induces apoptosis^12^. Nalm6 cells expressing Click Beetle Green luciferase (CBG) were edited to disrupt either *FADD* or *BID* (**Extended Data Figures 2a-b**). WT Nalm6 cells were combined with varying percentages of knockout (KO) Nalm6 cells (80%, 40% or 20% KO) and then combined with CART19. Co-cultures with higher fractions of KO cells were more resistant to death at 24h (**Extended Data Figures 3a-b**) and over time (**Figure 1e**), and demonstrated a progressive enrichment of KO cells during co-culture (**Extended Data Figures 3c-d**). Notably, disruption of *FADD* or *BID* did not protect Nalm6 from chemotherapy-mediated killing (**Extended Data Figure 4a**). Resistance was not unique to CD19-targeted CAR therapy, as we observed similar results using CD22-targeted CAR T cells (CART22, **Extended Data Figures 4b-d**). To establish that death receptor activity was essential for CAR T cell cytotoxicity, we disrupted the genes encoding the death receptor ligands Fas ligand (FasLG) or TRAIL in CART19 cells and combined them with WT Nalm6. Loss of FasL resulted in modest impairment in tumor killing, while loss of TRAIL caused near complete abrogation of CART19 cytotoxic function (**Extended Data Figure 4e**). To interrogate the role of death receptor signaling on CART19 activity *in vivo*, we established systemic disease in NOD/SCID/γc^-/-^ (NSG) mice^13^ with WT, *FADD*^KO^ or *BID*^KO^ Nalm6, and then delivered control (un-engineered) T cells or a sub-curative dose of CART19. Animals with WT tumors had an anticipated significant but transient disease response, while animals with KO tumors demonstrated rapid disease progression despite CART19 treatment (**Extended Data Figures 4g-h**). Remarkably, CART19 treatment led to a marginal improvement in median survival (4-5 days) in animals with KO tumors (**Figures 1f-g**), demonstrating that disruption of a single death receptor signaling gene can lead to rapid CART19 failure *in vivo*. These data indicate that disruption of death receptor signaling in ALL enables specific resistance to CAR T cells, and that death receptor engagement is required for CART19 activity.

T cells can kill target cells either by ligation of surface death receptors to initiate the extrinsic apoptotic pathway, or by secretion of cytolytic molecules (ie. perforin and granzyme) that activate the intrinsic apoptotic pathway^12^. Interestingly, resistance to extrinsic apoptosis was not overcome by initiation of intrinsic apoptosis in our *in vitro* and *in vivo* studies. To investigate this, we evaluated how impairment of tumor death receptor signaling impacted other CART19 functions. Consistent with our luciferase-based cytotoxicity assays (**Figures 1e-g**), evaluation of CART19 and Nalm6 co-cultures by flow cytometry demonstrated elimination of WT Nalm6 but only a transient decline in BID^KO^ Nalm6 burden (**Extended Data Figure 5a**). Following this diminished cytotoxicity, co-culture with *BID*^KO^ Nalm6 resulted in impaired CART19 expansion (**Figure 2a**), cytolytic molecule production and cytokine secretion over time (*Extended Data Figures 5b-f*), suggesting that failure to induce extrinsic apoptosis in target cells may impair CART19 function. To explore this hypothesis, we investigated the kinetics of this apparent acquired dysfunction. We sorted CART19 cells from short (5 days) or long (15 days) co-cultures with either WT or *BID*^KO^ Nalm6, and evaluated their ability to initiate effector functions upon reexposure to WT Nalm6 (**Figure 2b**). CART19 cells collected after short initial exposures to either target cell type demonstrated similar expansion, cytokine production and anti-tumor cytotoxicity upon re-exposure (**Figures 2c-e, Extended Data Figures 5g-j**). Long initial exposures, however, resulted in divergent functional abilities: CART19 cells cultured with BID^KO^ Nalm6 expanded poorly, produced less IFNγ, TNFα and MIP1α, and were unable to kill WT Nalm6 upon re-exposure (**Figures 2f-h, Extended Data Figures 5k-n**). Similar defects were induced when CART19 was continuously exposed to either *BID^KO^* or WT Nalm6 (**Extended Data Figure 6**), suggesting that dysfunction was not linked to impaired tumor death receptor signaling itself, but a result of the subsequent persistent antigen exposure. To further explore the role of persistent antigen exposure in a tumor-agnostic system, we combined WT Nalm6 with *TRAIL*^KO^ or perforin (*PRF1*)^KO^ CART19 cells, both of which have significantly impaired cytotoxic function (**Extended Data Figures 4e-f**). Consistent with our previous observations, *TRAIL*^KO^ and *PRF1*^KO^ cells demonstrated impaired expansion during initial exposure to WT Nalm6, and developed impairments in effector function during prolonged coculture (**Extended Data Figure 7**). Collectively, these data demonstrate that impaired death receptor signaling results in persistence of target antigen, which rapidly leads to CART19 dysfunction.

**Figure 2.**
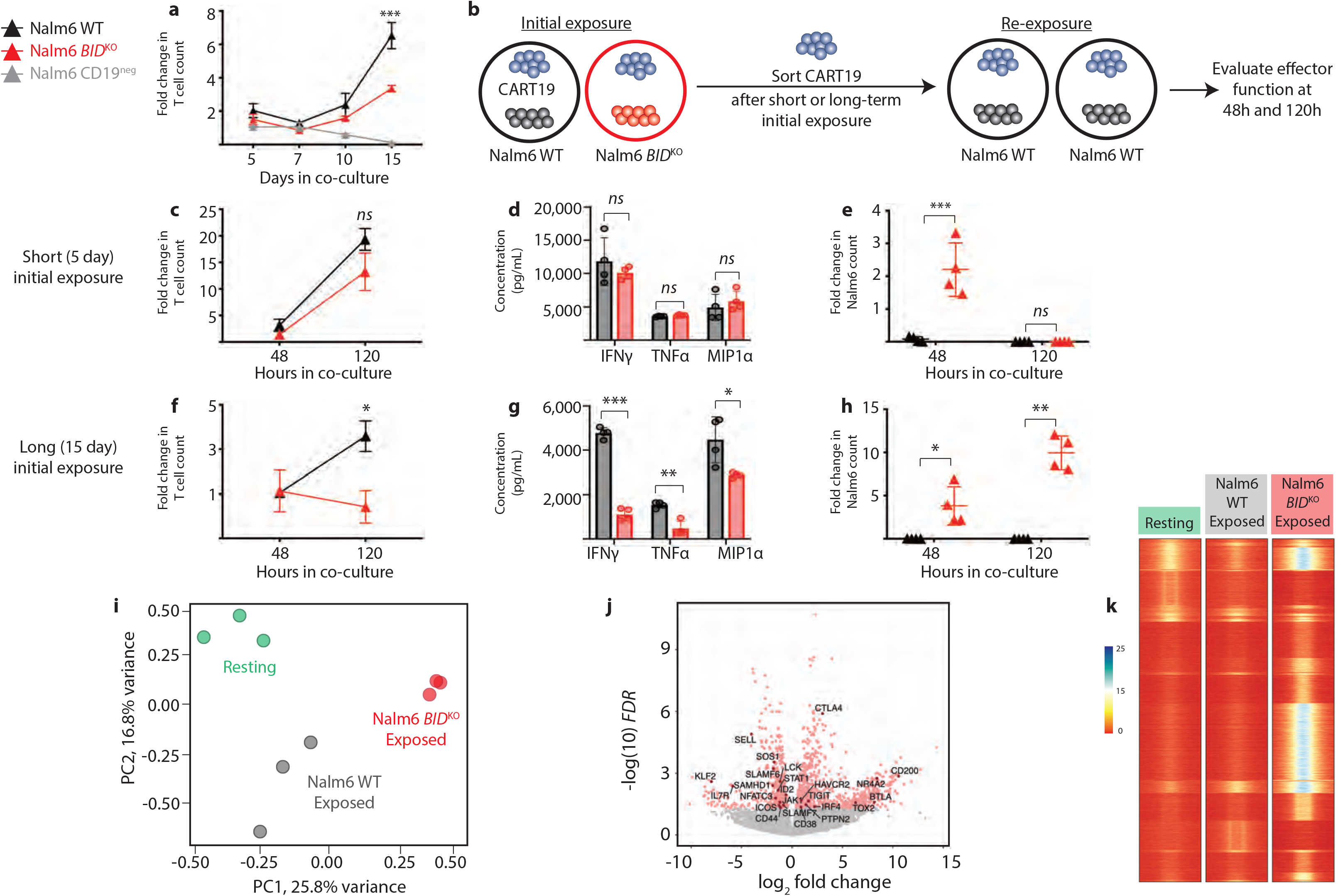
Exposure to death receptor-impaired ALL results in acquired CART19 dysfunction. **a,** WT, *BID*^KO^ or CD19^neg^ Nalm6 cells were co-cultured with CART19 cells and T cell expansion was measured over time. **b,** Schematic of functional evaluation studies, in which CART19 cells were sorted after short-term (5 day) and long-term (15 day) initial culture with either Nalm6 WT or *BID*^KO^ and then re-exposed to Nalm6 WT in secondary cultures. Measurement of effector functions including **c,f,** CART19 expansion after 48 and 120h of secondary culture, **d,g,** cytokine secretion after 48 hours, and **e,h,** Nalm6 survival after 48 and 120h of secondary culture. i, Principal component analysis of gene expression profiles CART19 cells at rest or after 15 days of exposure to Nalm6 WT or *BID*^KO^. **j,** Volcano plot of differentially expressed genes in CART19 cells after exposure to Nalm6 WT or *BID*^KO^. Transcripts with log-fold change >0.5 and FDR >0.05 are shown in red. **k,** Heatmap of differentially accessible chromatin sites in CART19 cells at rest or after exposure to WT or *BID*^KO^ Nalm6 *P<0.05, **P<0.01, ***P<0.001 by ANOVA

To explore the molecular programs underlying this T cell dysfunction, we performed phenotypic, transcriptomic and epigenomic interrogation of CART19 cells exposed to WT and *BID*^KO^ Nalm6. Persistent exposure to antigen is the central driver of T cell exhaustion in human and murine models of chronic viral infection and cancer^14–20^. Given the similarities in effector dysfunction (impaired expansion, cytokine production and cytotoxicity) between classically exhausted T cells and dysfunctional CART19 cells, we first evaluated expression of exhaustion-related immunosuppressive surface proteins PD-1 and Tim3. We observed a significant rise in expression of both markers on CART19 cells exposed to *BID*^KO^ Nalm6 or continuously exposed to WT Nalm6 (**Extended Data Figures 8a-f**). However, unlike the pattern seen in classical T cell exhaustion, CART19 PD-1 levels returned to normal during the course of continuous antigen exposure. We also found that chronic antigen exposure led to persistent elevations in expression of the exhaustion-related transcription factor Tbet (**Extended Data figures 8g-h**). We next analyzed gene expression profiles of CART19 cells at rest or after 15 days of exposure to WT or *BID*^KO^Nalm6. These three groups clustered independently by principal component analysis (**Figure 2i**), and we identified 1285 significantly differentially expressed genes in CART19 cells exposed to WT or *BID*^KO^ Nalm6. GSEA of upregulated transcripts in *BID*^KO^ exposed CAR T cells demonstrated enrichment of cell cycle genes and metabolic regulators, accompanied by decreased expression of immunostimulatory genes (**Extended Data Figures 9a-b**). Specifically, CART19 cells exposed to *BID*^KO^ Nalm6 demonstrated increased expression of known exhaustion-related genes (*EGR2, CTLA4, IRF4, BTLA, TOX2*), as well as several other dysfunction-associated genes such as *NR4A2*, a central regulator in several types of T cell dysfunction^21,22^ (**Figure 2j**). Further, we found decreased expression of specific T cell activating genes, including TCR signaling molecules (*SLAMF6, LCK, JAK1, SOS1*), stimulatory surface receptors (*IL7R, ICOS*), and transcriptional activators (*STAT1, ID2, NFATC3*) in *BID*^KO^ exposed CART19. Together, these data reflect that exposure to *BID*^KO^ tumors drives a transcriptional program in CART19 that is concurrently highly metabolically active and highly immunosuppressed.

Given that CART19 dysfunction developed over time, we interrogated CART19 chromatin accessibility to determine if persistent antigen exposure results in epigenetic imprinting. Concurrently with RNA sequencing, we profiled CART19 cells using the assay for transposase-accessible chromatin with high-throughput sequencing (ATACseq). We observed significant chromatin remodeling in CART19 cells after exposure to *BID*^KO^ Nalm6, with more modest changes in CART19 cells exposed to WT Nalm6 (**Figure 2k**). Comparison of promoter site accessibility by GSEA revealed increased accessibility of immune activation genes and decreased accessibility of genes regulating metabolic processes and intracellular signaling in the *BID^KO^* exposed cells (**Extended Data Figures 9c-d**). These epigenetic patterns have been previously reported as an exhaustion-specific signature^14^. Despite these similarities to classically exhausted T cells, we did not observe increased chromatin accessibility at key exhaustion-related loci (*PDCD1, CTLA4, TIGIT, EOMES, CD38* or *PTGER2*, **Supplementary Table 2**)^14,16,20,23^. The foundational work describing the epigenetic features of exhaustion has explored chromatin modifications in endogenous T cells persistently stimulated through native T cell receptors and co-stimulatory molecules. Intriguingly, previous studies have shown that development of exhaustion in CAR T cells can be mitigated by replacing the CD28 co-stimulatory domain with the 41BB signaling domain^24,25^, suggesting that persistent stimulation of 4-1BB-bearing CARs may drive distinct cellular programs. To identify the unique changes associated with CART19 dysfunction resulting from persistent antigen exposure, we integrated our transcriptomic and epigenomic data to reveal genes with differential expression and concordant promoter accessibility changes. In *BID*^KO^ exposed CART19 cells, we found 41 genes with significantly increased expression and enhanced promoter accessibility as compared to WT exposed cells, several of which are key immunosuppressive transcription factors (*TOX2, IRF8, PRDM1*). Similarly, we found 14 genes with decreased expression and suppressed promoter accessibility, several of which are immunostimulatory (*DGKA, NMT2, TXNIP, THEMIS, KLF2*; **Supplementary Table 2**). Together, these findings suggest that exposure to leukemia cells that are deficient in death receptor signaling leads to global programmatic changes in CART19 that shares many features with classical T cell exhaustion,^14,16,20^ but also induces unique epigenetic changes in the setting of persistent CAR engagement.

In light of these observations, we investigated the association between tumor expression of death receptor genes and clinical outcomes after treatment with tisagenlecleucel, a clinical CD19-directed CAR T cell product^1^. We evaluated available leukemia-infiltrated bone marrow samples collected prior to treatment from patients who had either durable (>1 year) complete remissions (CR, *n* = 17) or no response (NR, *n* = 8) from two multi-center clinical trials of tisagenlecleucel for relapsed pediatric ALL (ClinicalTrials.gov identifiers NCT02435849 and NCT02228096). Using the KEGG pathway database, we identified the pro-apoptotic genes specifically involved in either intrinsic apoptosis or death receptor signaling to generate two distinct gene sets for analysis (**Supplementary Table 1** and **Methods**). Using normalized expression data from our patient samples, we generated two gene set scores (“intrinsic apoptosis signature” and “death receptor signature”) that represented overall expression of pro-apoptotic genes in each pathway. We found no difference in expression of the intrinsic apoptosis signature between CRs and NRs (**Extended Data Figures 10a-b**). In contrast, non-responders had lower expression of death receptor genes and significantly lower death receptor signature scores than durable responders (P=0.0126, **Figure 3a-b**). Using receiver operative curve analysis, we identified a gene set score cutoff that provided high positive (85.7%, 95% CI [70.6, 94.4]) and negative predictive values (88.9%, 95% CI [53.4, 98.2]) of durable remission after tisagenlecleucel (**Figure 3b**).

**Figure 3.**
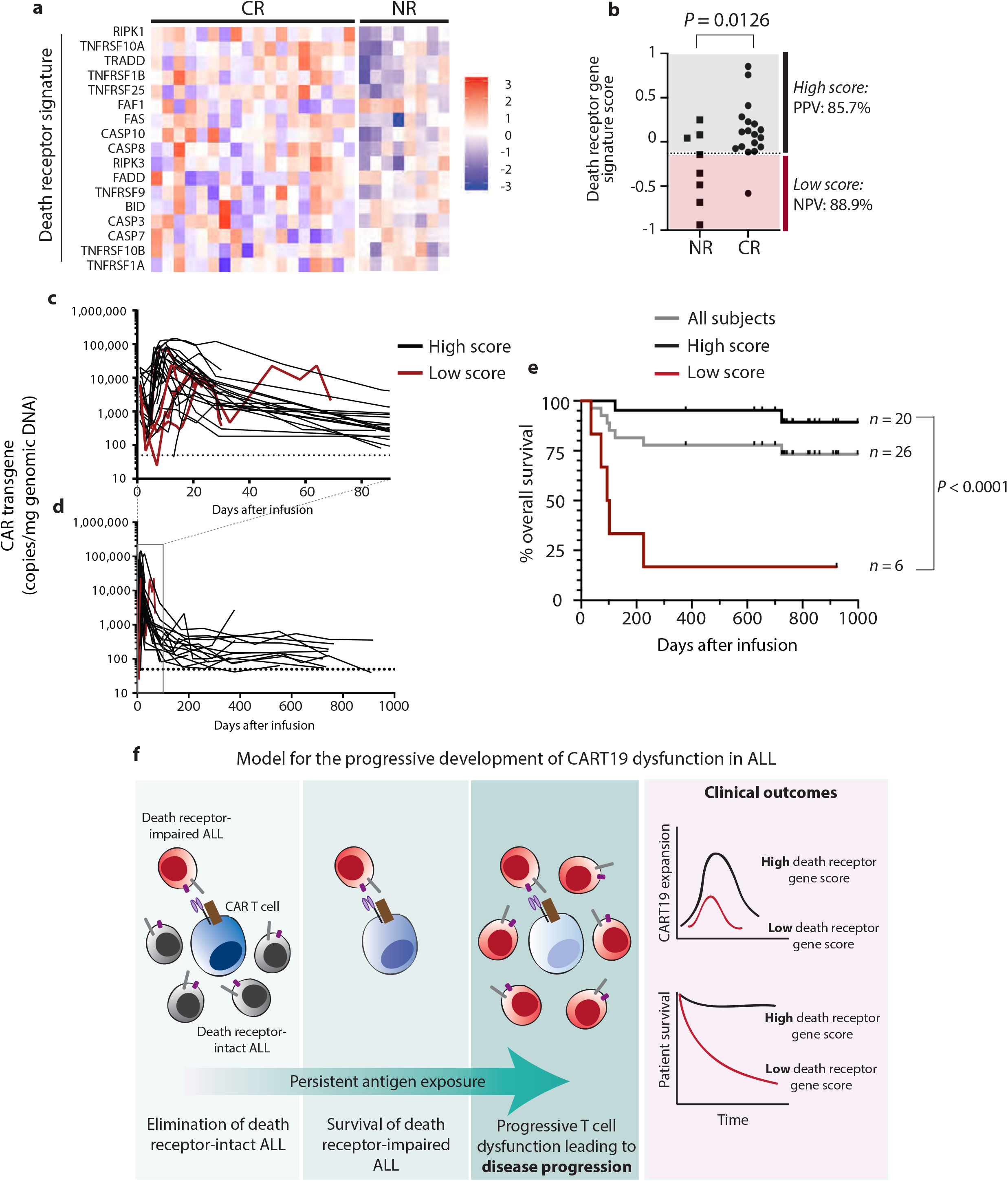
Death receptor gene expression correlates with outcomes after tisagenlecleucel. **a,** RNA expression heat map of pre-treatment leukemia-infiltrated bone marrow collected from patients treated with tisagenlecleucel (CR, *n*=18; NR, *n*=8). **b,** Integrated gene set expression score of death receptor-associated pro-apoptotic genes (“death receptor gene signature”) in CRs and NRs. **c-d,** Pharmacokinetic analysis of tisagenlecleucel in peripheral blood over the first **c,** 90 and **d,** 1000 days after infusion, as measured by quantitative PCR of CAR transcripts in patients with high (*n*=20) or low (*n*=6) death receptor signature scores. e, Overall survival of all patients analyzed. **f,** Proposed model for mechanism of impaired tumor cell death receptor signaling that leads to CART19 dysfunction and therapeutic failure.

Tisagenlecleucel expansion and persistence are highly associated with clinical response^26,27^ and based on our pre-clinical observations we hypothesized that exposure to death receptor-impaired leukemia may correlate with dysfunctional T cell expansion and persistence in patients. Pharmacokinetic analysis revealed reduced tisagenlecleucel expansion and shorter persistence in patients with death receptor signature scores below the cutoff (low scores) as compared to patients with scores above the cutoff (high scores) (**Figure 3c-d**). We further explored clinical outcomes in this patient cohort and found striking differences in overall survival between patients with high and low scores (**Figure 3e**, *P*<0.0001). Notably, 5 of 6 patients with low scores died soon after therapy. Given that clinical samples were not collected in a manner that allowed analysis of purified leukemic blasts, we adjusted gene set scores for the extent of blast involvement in patient marrow (see Methods) and repeated this analysis. These adjusted gene set scores were also highly-associated with clinical outcome, tisagenlecleucel expansion and persistence, and overall patient survival (**Extended Data Figure 10c-f**). Multivariate logistic regression analysis identified that the association between death receptor signature score and clinical response was highly mediated by the association between death receptor signature score and T cell expansion, mirroring our *in vitro* observations of impaired T cell expansion upon exposure to death receptor-impaired Nalm6.

Until now, primary resistance to CART19 was presumed to be a result of intrinsic T cell defects^28,29^, leading to extensive efforts to use heathy donor (i.e. allogeneic donor or “universal” donor) T cells as a substrate for CAR T cell manufacturing. Our data offer a model in which tumor-intrinsic biology contributes to observed therapeutic failure in a manner that is orthogonal to chemotherapy resistance (**Figure 3f**). These data suggest that antigen persistence resulting from slow response kinetics may also lead to progressive T cell dysfunction, tilting the balance in favor of primary resistance to CART19. Additional studies investigating the role of death receptor gene expression as a clinical biomarker may allow improved patient selection for CAR therapy products and trials. Further, development of methods to overcome death receptor-mediated resistance may significantly improve clinical outcomes after engineered cell therapy.

## Methods

### Brunello guide library screens

The Brunello sgRNA knockout plasmid library was designed and produced as previously described^6–7^. For generation of the lentiviral knockout library, four 15cm dishes of HEK293T cells (ATCC ACS-4500) were seeded at ~40% confluence the day before transfection in D10 media (DMEM supplemented with 10% fetal bovine serum). One hour prior to transfection, media was removed and 13mL of pre-warmed reduced serum OptiMEM media was added to each flask. For each dish, 20μg of plasmid library, 10μg of pVSVg, and 15μg of psPAX2 (Addgene) were diluted in 2mL OptiMEM. 100μL of Lipofectamine 2000 (Invitrogen) was diluted in 2mL OptiMEM and after a 5min incubation was added to the diluted DNA mixture. The complete mixture was incubated for 20min before being added to cells. After 6h, the media was changed to 15mL D10. Media containing lentiviral particles was removed after 48h and stored in 1mL aliquots at −80°C. To find optimal virus volumes to achieve an MOI of 0.2-0.3, each viral production was individually titrated. Briefly, 2×10^6^ cells were plated into each well of a 12 well plate, and varying volumes of virus (between 0 and 160μL) were added. Cells were counted and each well was split into two plates and diluted 1:10; one plate underwent selection with puromycin (1μg/mL). After 48h, each well was counted and viability assessed to determine percent transduction (calculated as cell count from the replicate with puromycin divided by cell count from the replicate without puromycin). The virus volume yielding a MOI closest to 0.2 was chosen for large-scale screening. For long-term depletion screening, 2×10^8^ Nalm6 cells (edited as described above) were combined with 5×10^7^ CART19 cells or control T cells (effector:target ratio 1:4) and co-cultured in standard culture media; 5×10^7^ control Nalm6 cells were frozen for genomic DNA analysis. Each condition was established in duplicate, and co-cultures were monitored by flow cytometry over time and maintained at a concentration of 1×10^6^ cells/mL. Cultures were monitored until Nalm6 recurred in selection cultures, at which time dead cells were removed using a Dead Cell Removal magnetic bead kit (Miltenyi) and living cells were frozen for genomic DNA analysis. Short-term screening co-cultures were established in the same manner, however dead cell removal and live cell collection was performed after 24h of co-culture.

### Genomic DNA extraction and guide sequencing

From each screening culture, 3×10^7^ −5×10^7^ were flash frozen as dry cell pellets. At time of DNA extraction, 6 mL of NK Lysis Buffer (50 mM Tris, 50 mM EDTA, 1% SDS, pH 8) and 30μL of 20mg/mL Proteinase K were added to the frozen cell sample and incubated at 55°C overnight. The following day, 30μL of 10mg/mL RNase A, diluted in NK Lysis Buffer to 10mg/mL and then stored at 4°C, was added to the lysed sample, which was then inverted 25 times and incubated at 37°C for 30min. Samples were cooled on ice before addition of 2mL of pre-chilled 7.5M ammonium acetate to precipitate proteins. The samples were vortexed at high speed for 20s and then centrifuged at ≥ 4,000×g for 10min. After the spin, a tight pellet was visible in each tube and the supernatant was carefully decanted into a new 15mL tube. 6mL 100% isopropanol was added to the tube, inverted 50 times and centrifuged at ≥ 4,000×g for 10min. Genomic DNA (gDNA) was visible as a small white pellet in each tube. The supernatant was discarded, 6mL of freshly prepared 70% ethanol was added to the tube and inverted 10 times and centrifuged at ≥4,000×g for 1min. The supernatant was discarded by pouring; the tube was briefly spun, and remaining ethanol was removed using a pipette. After air drying for 10-30min, 500μL of 1x TE buffer was added, the tube was incubated at 65°C for 1h and at room temperature overnight to fully resuspend the DNA. The next day, the gDNA samples were vortexed briefly, and gDNA concentration was measured.

To measure the distribution of sgRNA within each screen arm, we used Illumina Next Generation Sequencing applied to an amplicon generated from a single targeted PCR of the integrated sgRNA cassette^6^. Briefly, we used all collected gDNA (1000x coverage) divided into 100μL PCR reactions with 5μg of DNA per reaction. We used Takara ExTaq DNA Polymerase and the default mix protocol with the following PCR program: (95° 2min, (98° 10sec, 60° 30sec, 72° 30sec) x 24, 72° 5min). PCR products were gel purified using the QiaQuick gel extraction kit (Qiagen). The purified, pooled library was then sequenced on a HiSeq4000 with ~5% PhiX added to the sequencing lane. Quality assessment was done by qubit (for concentration), bioAnalyzer (for size distribution) and Kapa Library Quantification (for clusterable molarity).

### Genome-wide screening data analysis

To count the number of reads associated with each sgRNA in each Fastq file, we first extracted the sgRNA targeting sequencing using a regular expression containing the three nucleotides flanking each side of the sgRNA 20bp target. sgRNA spacer sequences were then aligned to a pre-indexed Brunello library (Addgene) using the short-read aligner ‘bowtie’ using parameters (-v 0 -m 1). Data analysis was performed using custom R scripts. Using the guide sequencing data, we analyzed enrichment and depletion of guide RNAs with the Model-based Analysis of Genome-wide CRISPR/Cas9 Knockout (MAGeCK) algorithm using the maximum likelihood estimation (MLE) module^32,33^. In our long-term screen, we found that over 25% of all sequenced reads in the experimental (CART19) selection condition originated from sgRNAs targeting CD19. This led to a highly skewed sgRNA distribution in the sequencing data, preventing meaningful statistical analysis of other targets identified using the MAGeCK algorithm.

### CRISPR/Cas9-guide design, genomic engineering and indel detection

CRISPR sgRNAs were designed using software integrated into Benchling (http://Benchling.com). For each target gene, six sgRNA sequences were designed to target early exon sequences, and *in vitro* transcribed using the GeneArt Precision gRNA Synthesis Kit (Invitrogen) for screening. Cells were electroporated using the Lonza 4D-Nucleofector Core/X Unit. Nalm6 cells were electroporated using the SF Cell Line 4-D Nucleofector Kit, and primary T cells were electroporated using the P3 Primary Cell 4-D Kit (Lonza). For Cas9 and sgRNA delivery, the ribonucleoprotein (RNP) complex was first formed by incubating 10μg of TrueCut Cas9 Protein V2 (Lonza) with 5μg of sgRNA for 10-30min at room temperature. Cells were spun down at 300 xg for 10min and resuspended at a concentration of 3-5×10^6^ cells/100μL in the specified buffer. The RNP complex, 100μL of resuspended cells, and 4μL of 100μM IDT Electroporation Enhancer (IDT) were combined and electroporated. Pulse codes EO-115 and CV-104 were used for primary T cells and Nalm6 cells, respectively. After electroporation, the cells were incubated in standard media at a 5×10^6^ cells/mL at 30°C for 48h, then cultured at 37°C for the duration of experimental procedures. TIDE (Tracking of Indels by DEcomposition) analysis was used to detect knock out (KO) efficiency at the genomic level^32^. Genomic DNA from electroporated cells was isolated (Qiagen DNeasy Blood & Tissue Kit) and 200-300ng were PCR amplified using Accuprime Pfx SuperMix and 10μM forward/reverse primers flanking the region of interest. Primers were designed such that the amplicon was at a target size ~1 kb. PCR products were gel purified and sequenced, and trace files were analyzed using Desktop Genetics software (tide.deskgen.com, Desktop Genetics) to determine KO efficiency. R^2^ values were calculated, reflecting goodness of fit after non-negative linear modeling by TIDE software^34^. See Supplemental Table 3 for sgRNA sequences and sequencing primer sequences.

### General cell culture

Unless otherwise specified, cells were grown and cultured at a concentration of 1×10^6^ cells/mL of standard culture media (RPMI 1640 + 10% FCS, 1% penicillin/streptomycin, 1% HEPES, 1% non-essential amino acids) at 37°C in 5% ambient CO2. All co-culture studies were performed an effector cell to target cell ratio of 1:4, unless otherwise stated.

### Lentiviral vector production and transduction of CAR-engineered human T cells

Replication-defective, third-generation lentiviral vectors were produced using HEK293T cells (ATCC ACS-4500). Approximately 8×10^6^ cells were plated in T150 culture vessels in standard culture media and incubated overnight at 37°C. 18-24h later, cells were transfected using a combination of Lipofectamine 2000 (96μL, Invitrogen), pMDG.1 (7μg), pRSV.rev (18μg), pMDLg/p.RRE (18μg) packaging plasmids and 15μg of expression plasmid (either CD19 or CD22-targeted CAR). Lipofectamine and plasmid DNA was diluted in 4mL Opti-MEM media prior to transfer into lentiviral production flasks. At both 24 and 48h following transfection, culture media was isolated and concentrated using high-speed ultracentrifugation (25,000 xg for 2.5 hours). Human T cells were procured through the University of Pennsylvania Human Immunology Core. CD4 and CD8 cells were combined at a 1:1 ratio, and activated using CD3/CD28 stimulatory beads (Thermo-Fisher) at a ratio of 3 beads/cell and incubated at 37°C overnight. The following day, CAR lentiviral vectors were added to stimulatory cultures at an MOI between 4-6. Beads were removed on day 6 of stimulation, and cells were counted daily until growth kinetics and cell size demonstrated they had rested from stimulation.

### Bioluminescence-based cell survival assays

Click beetle green (CBG)-engineered cell survival was measured using bioluminescent quantification. D-luciferin potassium salt (Perkin-Elmer) was added to cell cultures (final concentration 15μg/mL) and incubated at 37°C for 10min. Bioluminescent signal was detected using a BioTek Synergy H4 imager, and signal was analyzed using BioTek Gen5 software. Percent specific lysis was calculated using a control of target cells without effectors.

### Flow cytometry

Cells were resuspended in FACS staining buffer (PBS + 3% fetal bovine serum) using the following antibodies: CD3 (clone OKT3, eBiosciences), CD4 (clone OKT4, eBiosciences), CD8 (clone SK1, BD), PD-1 (clone J105, eBiosciences), TIM3 (clones F38-2E2, BioLegend), Tbet (clone 4B10, eBiosciences). CD19 CAR was detected using an anti-idiotype antibody provided by Novartis Pharmaceuticals. All changes in overall tumor or T cell counts reflect changes in absolute cell counts, which were determined using CountBright absolute counting beads (ThermoFisher). Cell viability was established using Live/Dead Aqua fixable staining kit (ThermoFisher), and data were acquired on an LSRII Fortessa Cytometer (BD). All data analysis was performed using FlowJo 9.0 software (FlowJo, LLC).

### Cytokine and cytolytic molecule quantification

Human cytokine quantification was performed using a custom 31-plex Luminex panel (EMD Millipore) containing the following analytes: EGF, FGF-2, Eotaxin, sIL-2Ra, G-CSF, GM-CSF, IFN-α2, IFN-γ, IL-10, IL-12P40, IL-12P70, IL-13, IL-15, IL-17A, IL-1RA, HGF, IL-1β, CXCL9/MIG, IL-2, IL-4, IL-5, IL-6, IL-7, CXCL8/IL-8, CXCL10/IP-10, CCL2/MCP-1, CCL3/MIP-1α, CCL4/MIP-1β, RANTES, TNF-α, VEGF. Cell culture supernatants were flash frozen on dry ice, and thawed at time of cytokine analysis. Assays were established per manufacturer recommendations. Data were acquired on a FlexMAP 3D quantification instrument, and analysis was done using xPONENT software. Data quality was determined by ensuring the standard curve for each analyte had a 5P R^2^ value > 0.95 with or without minor fitting using xPONENT software. To pass assay technical quality control, the results for two controls in the kit were required to be within the 95% confidence interval provided by the vendor for >25 of the tested analytes. We established *in vitro* quantification of granzyme B (antibody sets from R&D Systems) and perforin (antibody sets from MABTECH), using ELISA substrate ADHP from Cayman Chemical, assay plates from E&K Scientific, and substrate plates from Greiner Bio-One. All ELISA reagents were prepared according to the protocols for DuoSet ELISA except for Color Reagent B, which was supplemented with ADHP at 100μM. Instead of coating the capture antibody to the wells of the ELISA plate, it was coated on the surfaces of macrospheres, which enabled the measurement of both analytes using 100μl of cell supernatant (collected as described above). Assays were set up using bead-strips in assay plates based on an assay map following the protocol for antibody sets. At the end of the assay, one substrate plate per 12 bead-strips for each analyte was prepared by adding 100μl/well of substrate solution (1:1 of color reagent A and ADHP). Each bead-strip was placed in one column of the substrate plate according to the assay map. Color development was for 10 to 30 mins. Plates were read on a FLUO STAR OMEGA instrument using fluorescent channel at 530 nm (excitation) and 590 nm (emission). Data were analyzed using Omega Data Analysis software.

### Xenograft mouse models

6-10 week old NOD-SCID-γc^-/-^ (NSG) mice were obtained from the Jackson Laboratory and maintained in pathogen-free conditions. Animals were injected via tail vein with 1×10^6^ WT, BID^KO^ or FADD^KO^ Nalm6 cells in 0.2mL sterile PBS. On day 7 after tumor delivery, 5×10^5^ T cells (control or CAR+) were injected via tail vein in 0.2mL sterile PBS. Animals were monitored for signs of disease progression and overt toxicity, such as xenogeneic graft-versus-host disease, as evidenced by >10% loss in body weight, loss of fur, diarrhea, conjunctivitis and disease-related hind limb paralysis. Disease burdens were monitored over time using a the Xenogen IVIS bioluminescent imaging system, as previously described^13^.

### RNA sequencing and analysis

RNA-seq libraries were made following the previously established SMARTseq2 protocol^35^. Briefly, total RNA was extracted using Qiazol (Qiagen) from 300 CART19 cells and recovered by RNA Clean and Concentrator spin columns (Zymo) followed by incubation with Oligo-dT. The transcription reaction was carried out on 100pg of cDNA for 1min at 55°C. Libraries were uniquely barcoded^36^ and amplified for 14 cycles. Fragment size distribution was verified, and paired-end sequencing (2 x 75 bp reads) was carried out on an Illumina NextSeq 500.

Raw reads were mapped to the GRCh37/hg19 genome assembly using the RNA-seq aligner STAR (version 2.5.4b)^37^. The algorithm was given known gene models provided by GENCODE (release_27_hg19, www.gencodegenes.org) to achieve higher mapping accuracy. Quantification was also performed by STAR using the --quantMode GeneCounts flag. Raw counts were normalized (TMM normalization implemented in edgeR, followed by the voom transformation). Lastly the Bioconductor package limma was used to assess differential expression using linear models. Genes with differential expression at a false discovery rate (FDR) < 0.05 and a log fold change (LFC) > 0.5 between replicates were considered for further analysis. Visualization was performed using the R package ggplot2.

### ATAC sequencing

Omni ATAC-seq libraries were made as previously described^38^. Briefly, nuclei were isolated from 50,000 sorted CART19 cells, followed by the transposition reaction using Tn5 transposase (Illumina) for 30 minutes at 37°C with 1000rp mixing. Purification of transposed DNA was completed with DNA Clean and Concentrator (Zymo) and fragments were barcoded with ATAC-seq indices^36^. Final libraries were double size selected using AMPure beads prior to sequencing. Paired-end sequencing (2 x 75 bp reads) was carried out on an Illumina NextSeq 500 platform. Adapters were trimmed using attack (version 0.1.5, https://atactk.readthedocs.io/en/latest/index.html), and raw reads were aligned to the GRCh37/hg19 genome using bowtie^39^ with the following flags: --chunkmbs 2000 --sam --best --strata -m1 -X 2000. MACS2^40^ was used for peak calling with an FDR cutoff of 0.05. Downstream analysis and visualization was done using HOMER^41^ and deepTools2^42^.

### Apoptotic pathway gene set development

Using the Kyoto Encyclopedia of Genes and Genomes (KEGG) “Apoptosis” pathway (hsa04210), we established two distinct gene set signatures. We determined to which pathway each gene belonged, either extrinsic (death receptor-dependent) or intrinsic (death receptor-independent) apoptosis. Genes that did not fall into either pathway (ie. nuclear factor kappa-light-chain-enhancer of activated B cells-associated, nerve-growth factor-associated)^43^ were excluded. Genes were then classified for activity, either pro-apoptotic or anti-apoptotic, and pro-apoptotic genes were included in gene sets.

### Evaluation of clinical specimens

Bone marrow (BM) specimens were collected from patients enrolled in two multi-center clinical trials (NCT02435849/ELIANA and NCT02228096/ENSIGN). All aspects of this study were approved by the Institutional Review Board at each participating institution. All relevant ethical regulations were complied with in this manuscript, and informed consent obtained at time of study enrollment allowed for RNAseq analysis on all patient samples evaluated. For purposes of this analysis, patients were categorized as achieving complete remission (CRs) or non-responders (NRs). CRs were defined as patients who achieved either complete remission or complete remission with incomplete hematologic recovery within 3 months and stayed in complete remission for at least one year after CAR infusion^1^. The cellular kinetics of tisagenlecleucel were determined as previously described^28^. Briefly, measurement of the synthetic CAR transgene by quantitative PCR. The cellular kinetic parameter of expansion, was derived from peripheral blood samples and estimated by noncompartmental methods using Phoenix (Pharsight, Version 6.4). Graphical analysis of transgene persistence was performed and categorized by high and low scores. Bone marrow blast percentage quantification for each patient was assessed by the Minimal Residual Disease (MRD) assay. Briefly, bone marrow aspirates were collected in sodium heparin vacutainer tubes, maintained at room temperature and tested within 5 days post collection. Immunophenotyping was performed starting with approximately 2×10^7^ total white blood cells (WBCs). WBC gate was adjusted to include WBCs and exclude red blood cells, dead/dying cells and debris. The number of viable WBCs identified by 7-AAD were used for calculating the MRD as a percent of total WBCs. A 4-tube, 8-color flow cytometry assay was performed using the BD FACSCanto II instrument to determine B-ALL MRD and Leukemia-associated Immunophenotypes (LAIP).

### RNA sequencing and bioinformatics analysis on clinical trial samples

Total RNA was extracted from bone marrow cells stored in PAXgene tubes according to the manufacturer’s instructions (Qiagen). Integrity was checked on the Agilent TapeStation (RIN), followed by preparation for sequencing using the TruSeq RNA v2 prep (Illumina). High-throughput sequencing was performed on an Illumina HiSeq 2500 platform to a target depth of 50 million paired-end reads per sample. Fastq files were processed for data QC, read mapping, transcript assembly, and transcript abundance estimation. A number of quality control metrics were assessed including data quality and GC content on per base and sequence levels, sequence length distribution and duplication levels, and insert size distribution. Finally, HTSeq was used to count the number of reads mapping to each gene^44^. Data normalization for differential expression analysis was carried out with the edgeR R package^45^. The logCPM value of each gene was z-normalized and gene set scores were calculated as the mean of the normalized gene expression for each gene in a given gene set. In order to account for the variability of the marrow blast count, the analysis was repeated and the gene set score was normalized to regress out the blast percentage. To identify a threshold gene set score with high positive and negative predictive value of clinical outcomes, we performed receiver operating curve (ROC) analysis on the cohort of gene set scores.

### Statistical analysis

All comparisons between two groups were performed using either a twotailed unpaired Student’s t-test or Mann-Whitney test, depending on normality of distribution. Comparisons between more than two groups were performed by two-way analysis of variance (ANOVA) with Bonferroni correction for multiple comparisons. All results are represented as mean ± standard error of the mean (s.e.m.). Survival data were analyzed using the Log-Rank (Mantel-Cox) test.

### Data availability

Guide library sequencing, RNA sequencing and ATAC sequencing data are available from Gene Expression Omnibus (GEO) using the accession number GSE130663 for all data sets. RNAseq and ATACseq expression data are included in Supplementary Table 2. Other data generated from this manuscript are available from the corresponding authors upon reasonable request.

## Supplementary Information

Supplementary Tables 1-3.

## Acknowledgements

The research was supported by the Society of Immunotherapy for Cancer Holbrook Kohrt Immunotherapy Translational Fellowship (N.S.), Breakthrough Bike Challenge Buz Cooper Scholarship (N.S.), NCI K08CA194256 (S.G.), American Society of Hematology Scholar Award, NCI 1K99CA212302, and R00CA212302 (M.R.), University of Pennsylvania-Novartis Alliance (S.G. and C.H.J.) and NCI 1P01CA214278 and R01CA226983 (C.H.J.). The authors thank A. Hoshino, S. Aggarwal, S. Kuramitsu, A. Green, and I. Maillard for valuable discussions and intellectual input, S. Lacey, F. Chen and N. Kengle for technical assistance with cytokine quantification assays, and J. Schug for informative discussions about guide library sequencing. The CD19 CAR used in these studies was generously provided by Dr. Dario Campana.

## Author contributions

N.S., C.H.J., S.G. and M.R. designed the research and wrote the manuscript. N.S., O. Shestova and P.R. performed *in vitro* screens, cell culture studies and animal experiments. O. Shalem generated the Brunello library. K.H. and M.D.W. analyzed sgRNA library, transcriptomic and epigenomic sequencing data. C.R.G. and S.L.B. performed transcriptomic and epigenomic library preparation and sequencing. E.J.O. and K.T.M. provided patient transcriptional and pharmacokinetic data. S.A.G. was the study steering committee chair for ELIANA and ENSIGN trials. N.S., E.J.O. and K.T.M. analyzed patient data. A.H. performed gene disruption and indel analyses. N.S. developed death receptor and intrinsic apoptotic gene signature sets. All authors reviewed the manuscript.

---

**Extended Data Figure 1.**
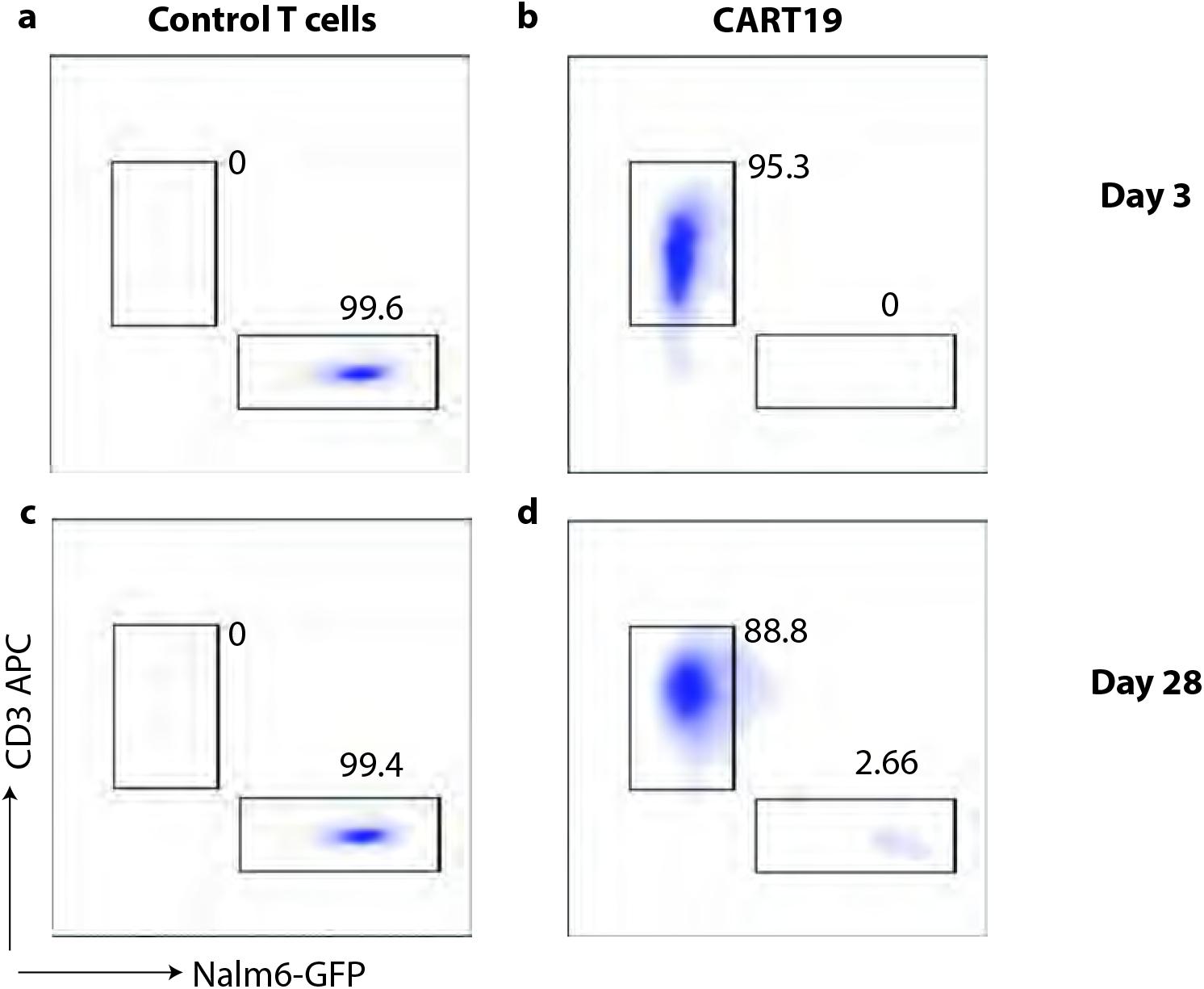
Brunello-edited ALL relapses under CART19 selective pressure *in vitro*. Representative flow cytometry plots from long-term relapse screen after **a-b,** 3 days of co-culture, and **c-d,** 28 days of co-culture with either control T cells or CART19.

**Extended Data Figure 2.**
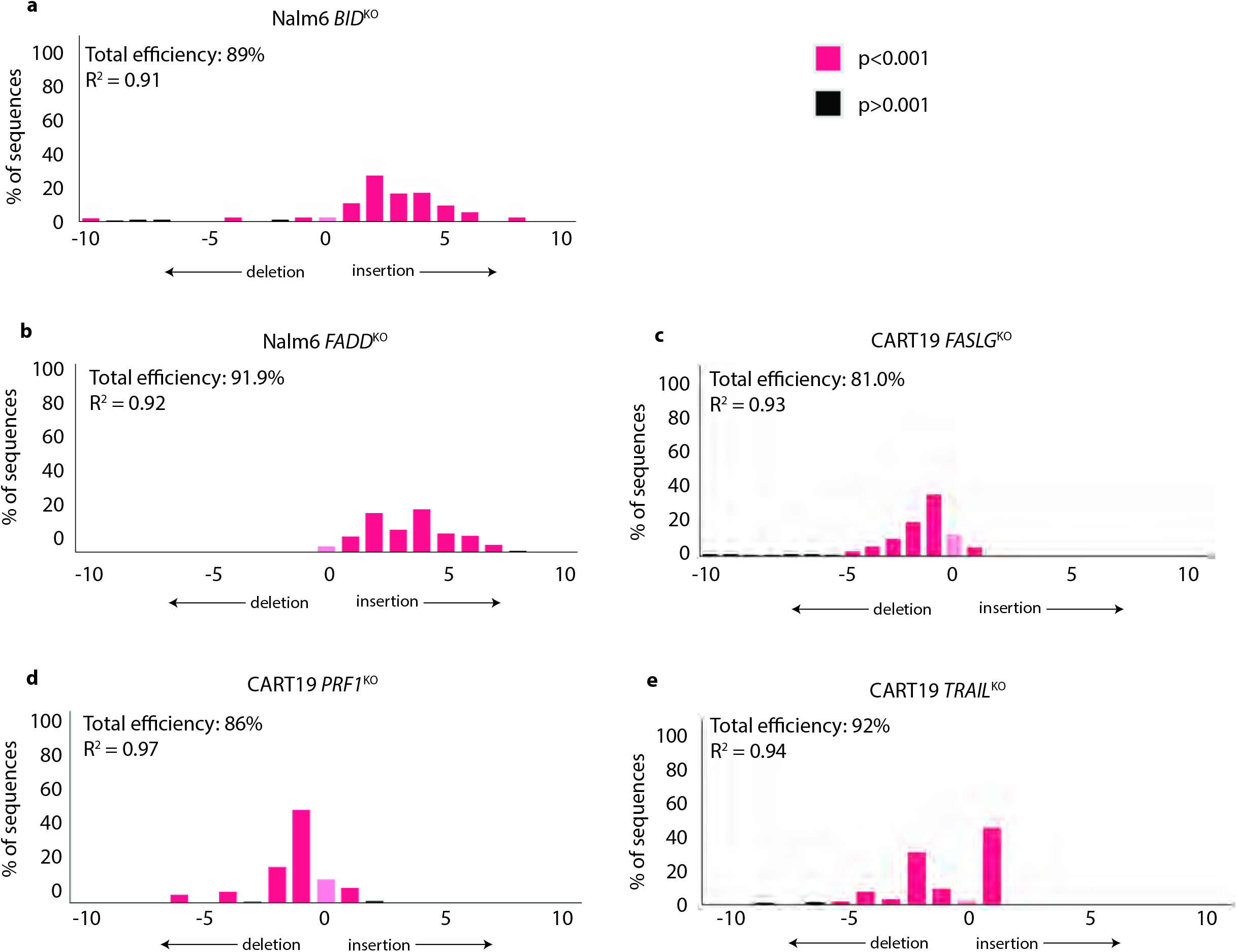
Targeted CRISPR/Cas9 editing results in high-efficiency of gene disruption in Nalm6 and CART19. **a-b,** Nalm6 ALL cells or **c-e,** primary human T cells engineered to express a CD19 CAR were edited using CRISPR/Cas9-directed sgRNAs, and gene disruption was evaluated using tracking of insertion and deletion (TIDE) sequencing^31^. Panels represent frequency of disruption at each upstream or downstream base relative to target disruption site.

**Extended Data Figure 3.**
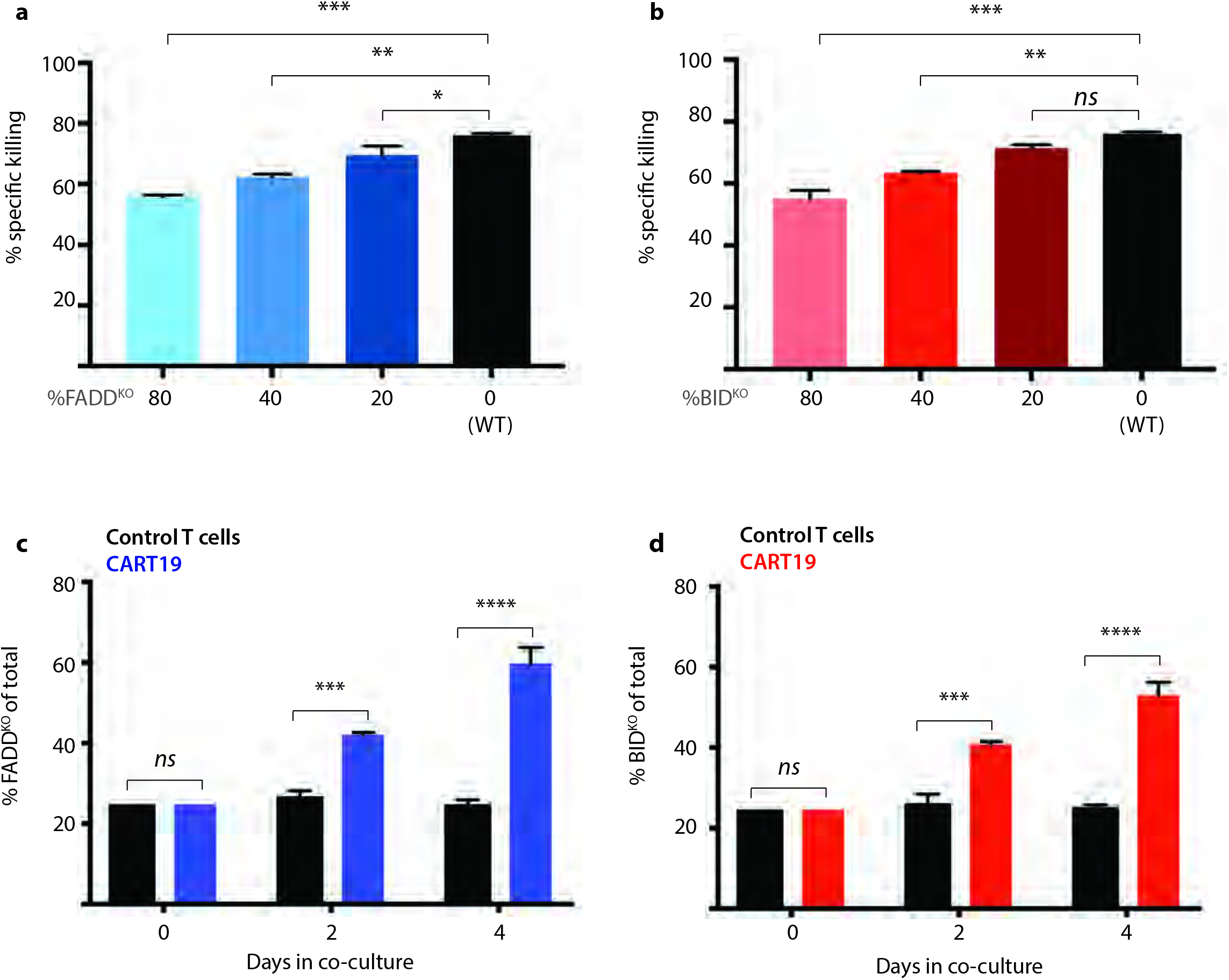
Loss of BID or FADD results in resistance to CART19. **a,** Nalm6 *FADD*^KO^ or **b,** Nalm6 *BID*^KO^ cells were combined with Nalm6 WT cells at progressive KO concentrations and co-cultured with CART19 and Nalm6 survival was measured at 24h. **c,** GFP+ Nalm6 *FADD*^KO^ or d, GFP+ Nalm6 *BID*^KO^ were combined with GFP-negative WT cells (25% GFP+KO with 75% GFP-neg WT cells) and co-cultured with either control T cell or CART19 cells. Proportion of GFP+ cells over time is shown. *P<0.05, **P<0.01, ***P<0.001, ****P<0.0001 by AN¤VA.

**Extended Data Figure 4.**
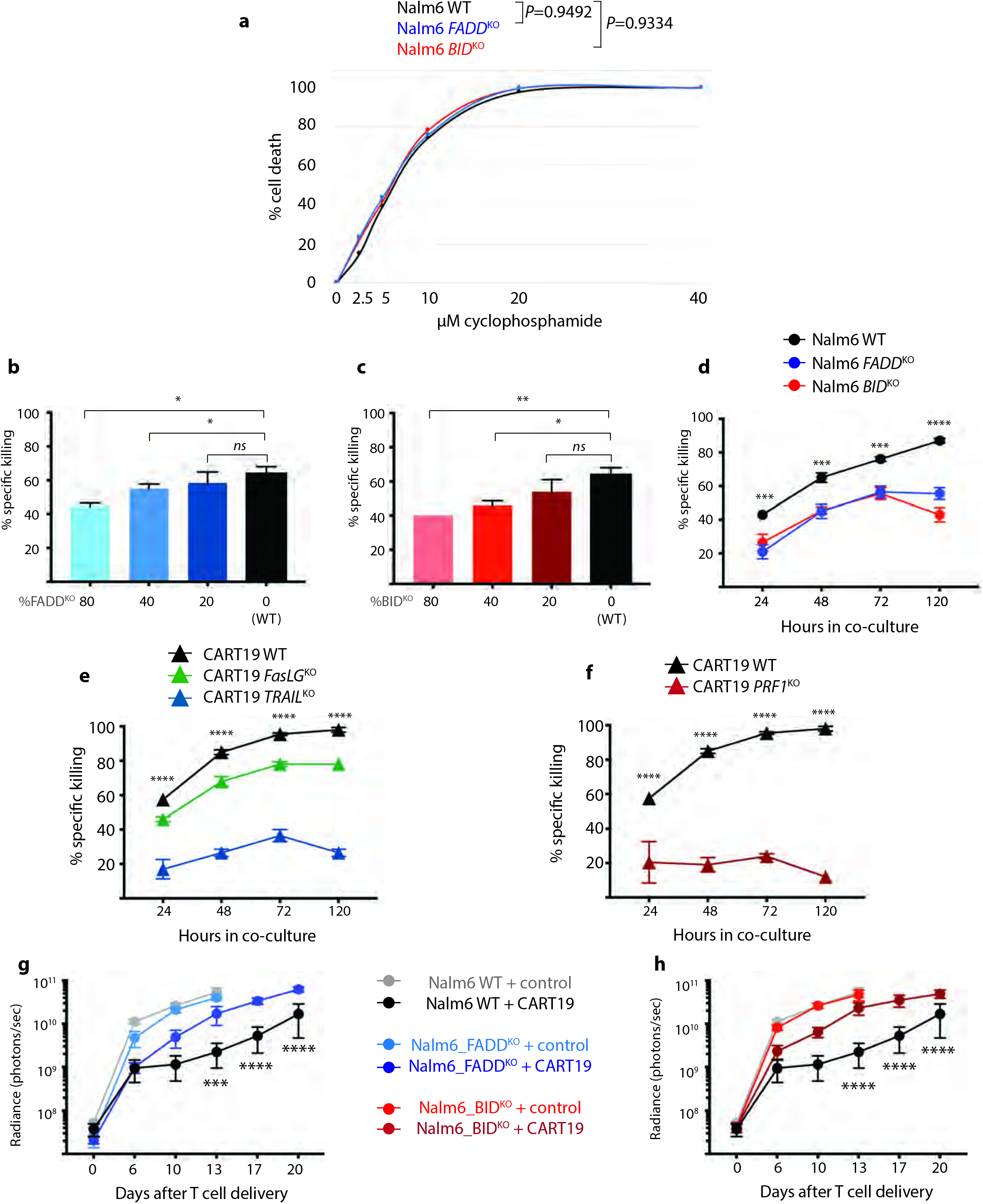
Impaired tumor death receptor signaling results in resistance to T cell killing. **a,** Nalm6 cells (WT, *FADD*^KO^ or *BID*^KO^) were exposed to increasing concentrations of cyclophosphamide and cell survival was measured at 24 hours. **b-d,** Nalm6 cells (WT, *FADD*^KO^ or *BID*^KO^) were co-cultured with CART22 cells as previously described (Figure 2) and Nalm6 survival was measured at **b-c,** 24h and **d,** over time. **e,** Survival of WT Nalm6 cells cultured with CART19 cells edited for loss of death receptor ligands. Significance reflects difference between WT and *TRAIL*^KO^ samples. **f,** Survival of WT Nalm6 cells cultured with perforin-deficient CART19. g-h, Disease progression over time in immunodeficient mice engrafted with luciferase-expressing Nalm6 (WT, *BID*^KO^ or *FADD*^KO^) after treatment with 5×10^5^ control T cells or CART19. *P<0.05, **P<0.01, ***P<0.001, ****P<0.0001 by ANOVA.

**Extended Data Figure 5.**
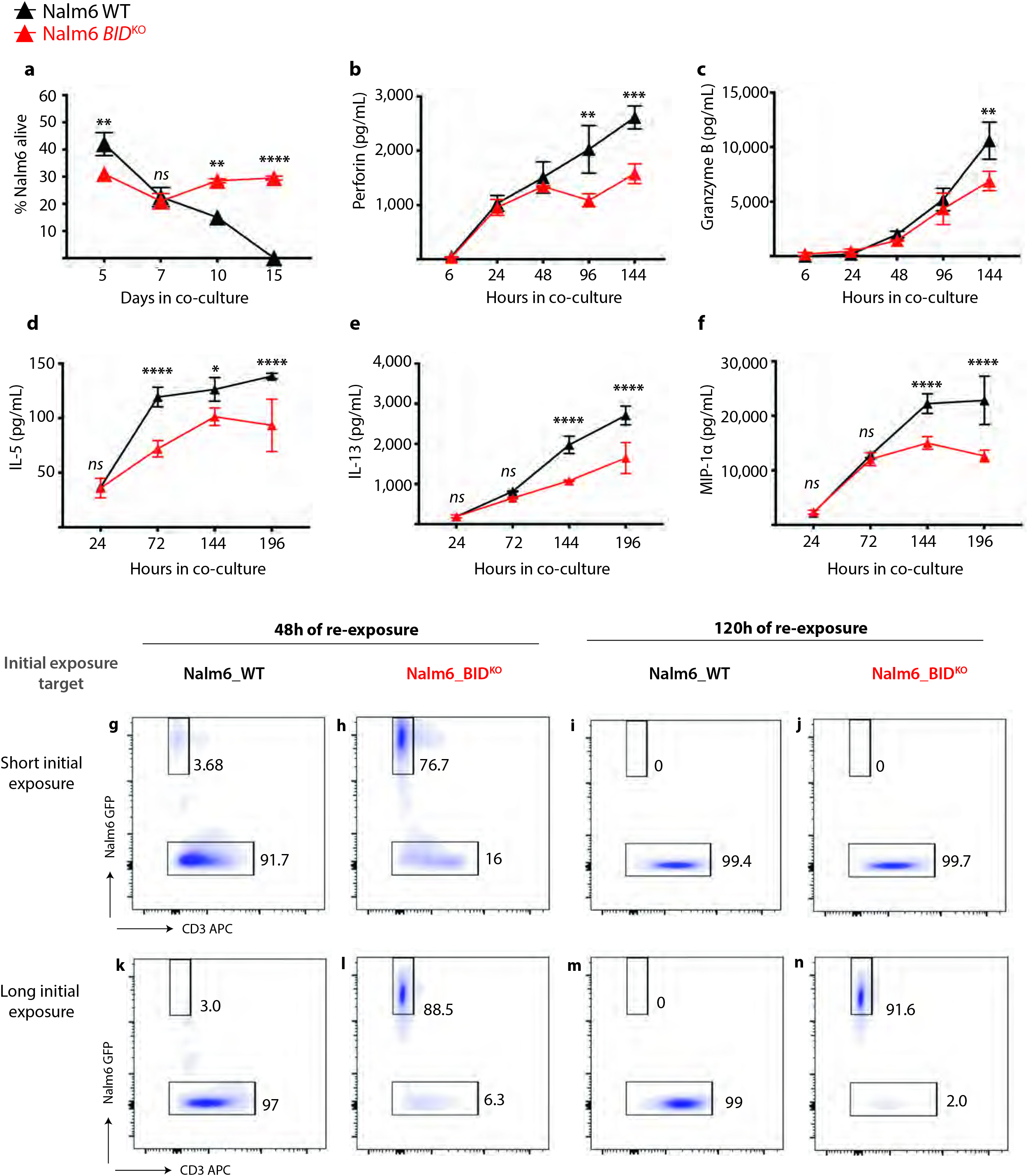
Exposure to death receptor impaired tumor cells results in progressive CART19 dysfunction. **a,** Nalm6 survival over time after treatment with CART19. **b-f,** CART19 cells were co-cultured with WT or *BID*^KO^ Nalm6 and supernatants were analyzed for **b,** perforin, **c,** granzyme B, **d,** IL-5, **e,** IL-13, or **f,** MIP1α. **g-n,** representative flow cytometry plots of sorted CART19 cells re-exposed to Nalm6 WT cells after **g-j,** short-term or **k-n,** long-term initial exposures.

**Extended Data Figure 6.**
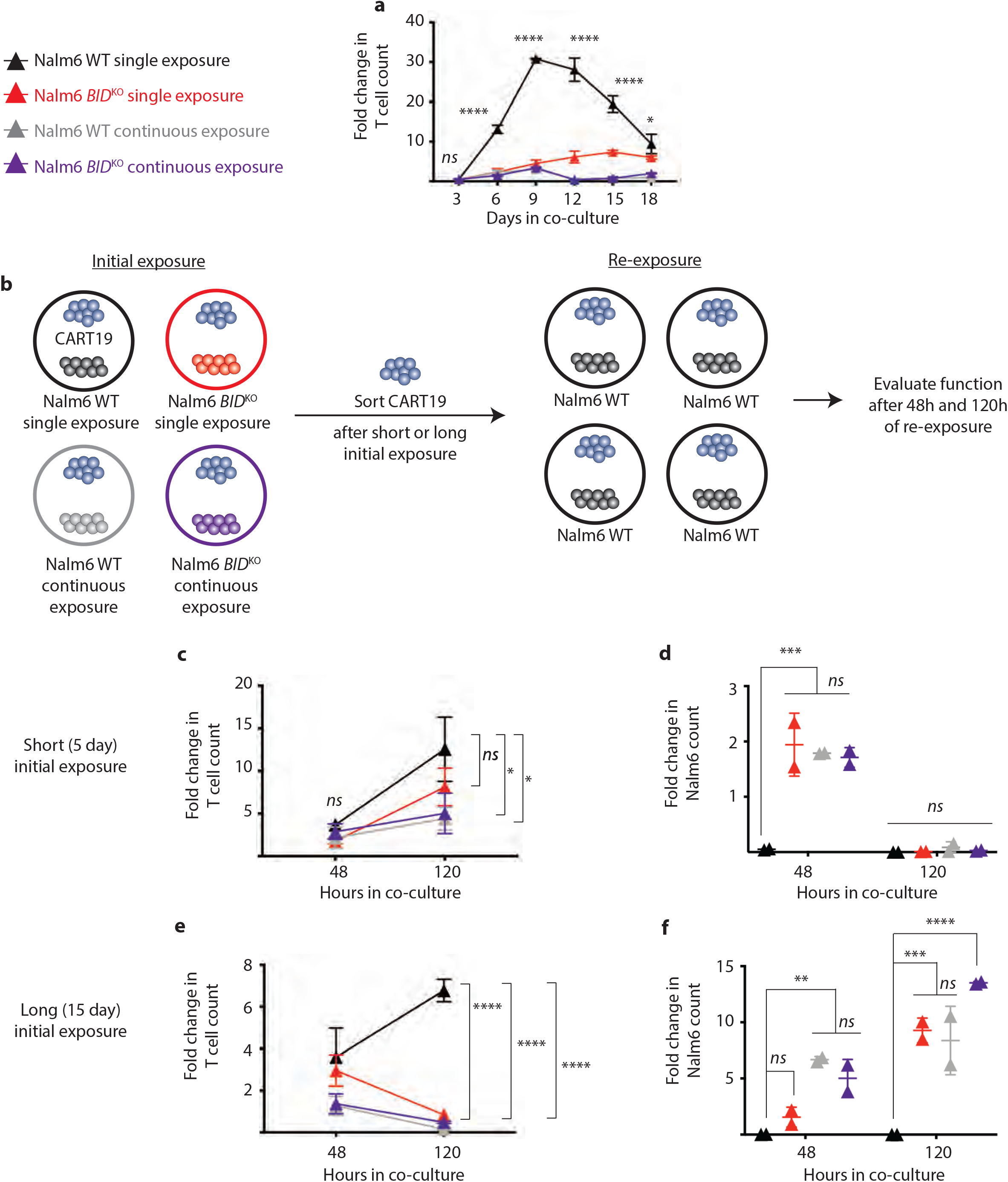
Persistent exposure to WT leukemia causes progressive CART19 dysfunction. CART19 cells were combined with Nalm6 WT or Nalm6 *BID*^KO^ once at the beginning of co-culture or repeatedly during co-culture to maintain a constant cell number. **a,** CART19 cells were quantified over time. **b,** Schematic of functional evaluation studies, as described in Figure 2. Measurement of **c,e,** CART19 expansion and **d,f,** Nalm6 WT survival over time. *P<0.05, **P<0.01, ***P<0.001, ****P<0.0001 by ANOVA.

**Extended Data Figure 7.**
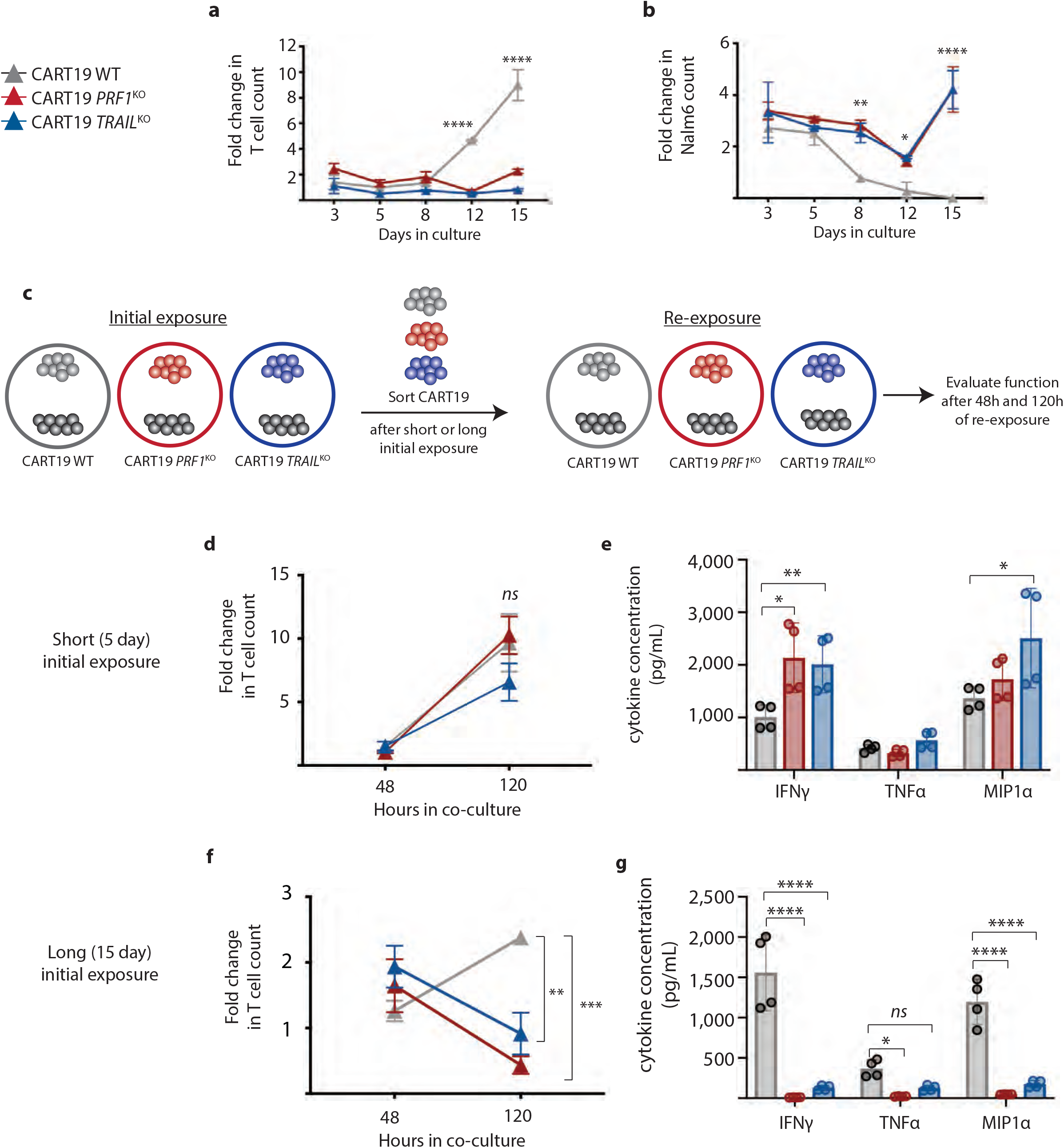
Persistent antigen exposure resulting from impaired T cell cytotoxicity machinery results in progressive CART19 dysfunction. **a-b,** WT, *TRAIL*^KO^ or *PRF1*^KO^ CART19 cells were combined with WT Nalm6 and **a,** CART19 expansion and **b,** Nalm6 survival were measured over time. **c,** Schematic of functional evaluation studies, in which CART19 cells (WT, *TRAIL*^KO^ or *PRF1*^KO^) were sorted after short-term (5 day) and long-term (15 day) initial culture with WT Nalm6, and then re-exposed to WT Nalm6. Measurement of **d,f,** CART19 expansion over time and e,g, effector cytokine secretion after 48 hours of re-exposure. *P<0.05, **P<0.01, ***P<0.001, ****P<0.0001 by ANOVA.

**Extended Data Figure 8.**
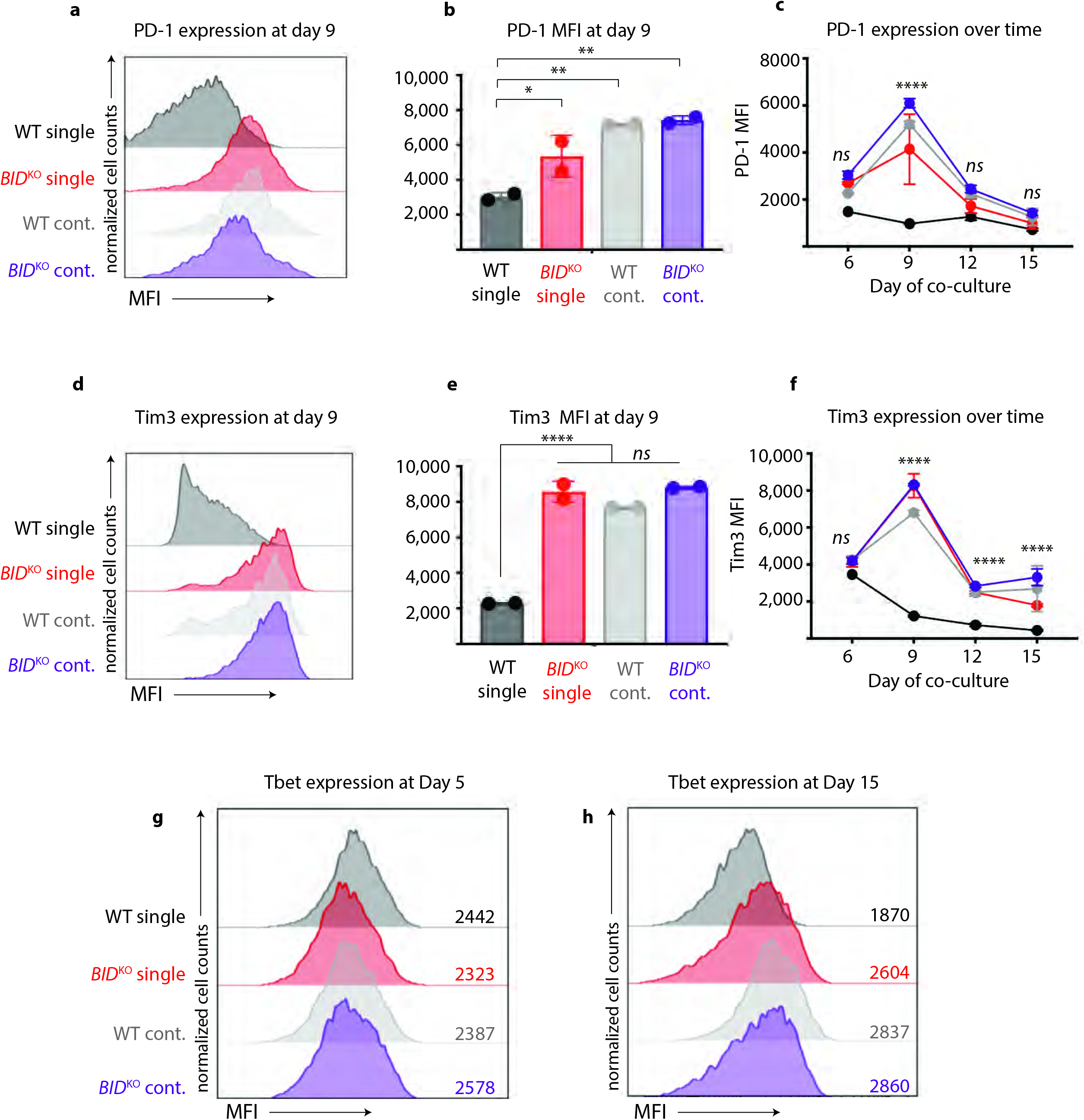
Persistent antigen exposure results in high inhibitory receptor and Tbet expression. **a-f,** CART19 expression of PD-1 and Tim3 was measured over time during co-culture with target Nalm6 cells. **a, d,** Representative flow cytometry plots and **b,e,** mean fluorescence intensity quantification at day 9 of co-culture and **c,f,** expression over time of co-culture. Tbet expression in CART19 cells at **g,** day 5 and **h,** day 15 of co-culture with WT or *BID*^KO^ Nalm6 after single or continuous exposure. *P<0.05, **P<0.01, ***P<0.001, ****P<0.0001 by ANOVA.

**Figure 9.**
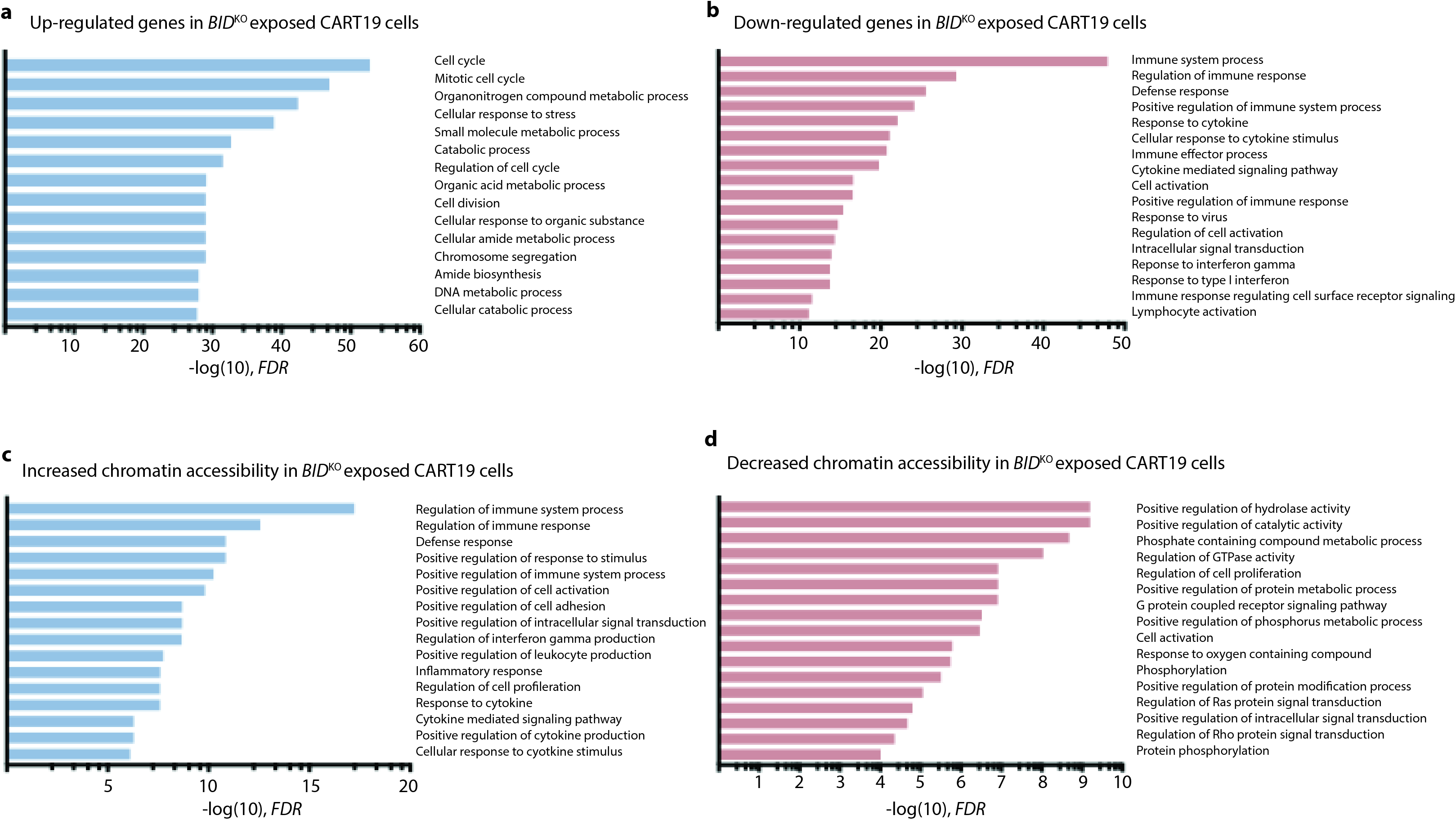
GSEA of transcriptomic and epigenomic sequencing of CART19 cells exposed to either WT or *BID*^KO^ Nalm6. **a-b,** sequenced transcripts or **c-d,** chromatin accessibility at promoter sites that were found to be significantly different (FDR<0.05, log-fold change > or <0.5) were analyzed using gene ontology (GO) pathway terms to identify pathways that were a, up-regulated, **b,** down-regulated, **c,** more accessible, or **d,** less accessible in *BID*^KO^ exposed CART19 cells.

**Extended Data Figure 10.**
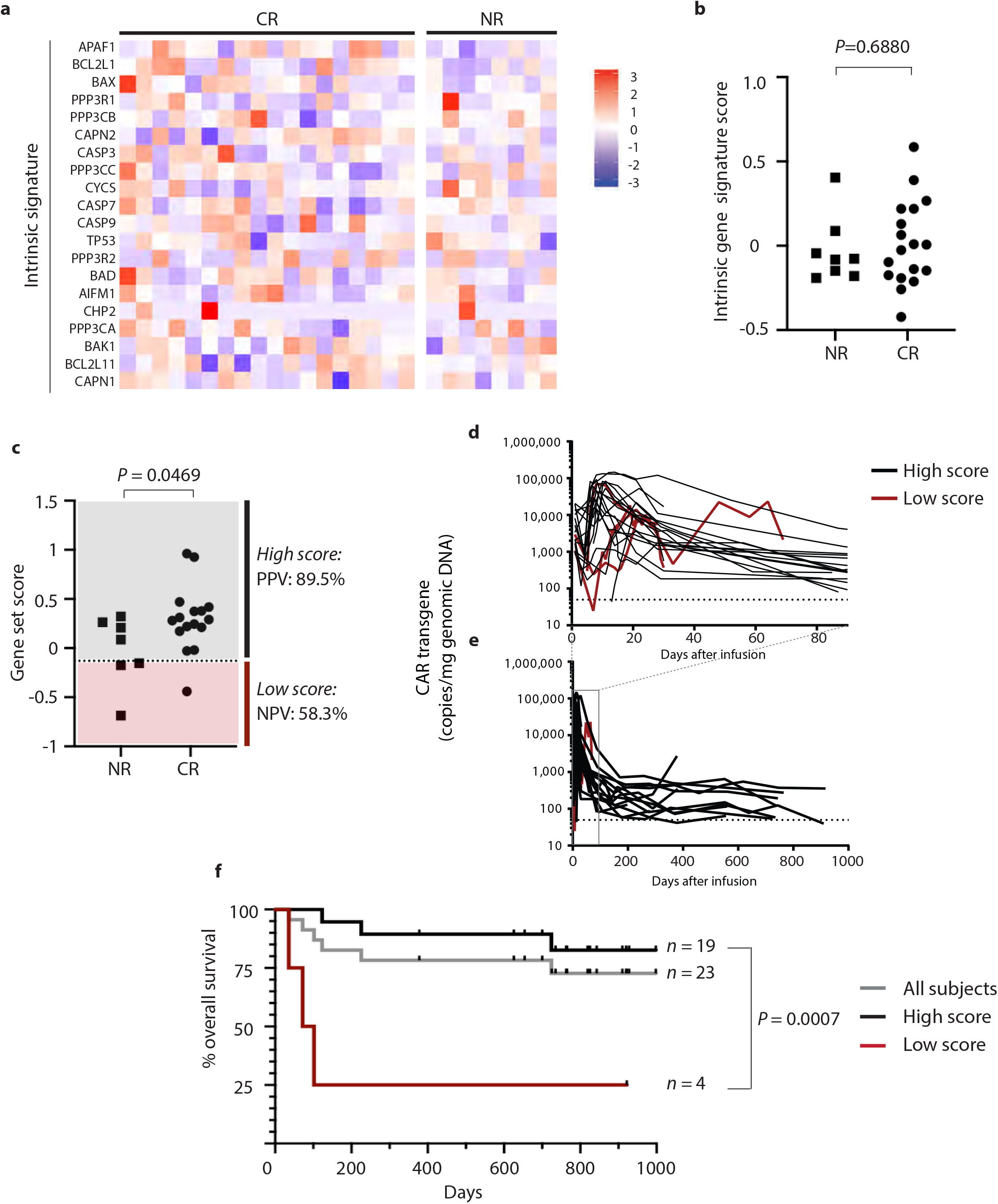
Death receptor gene expression signature is associated with clinical outcomes. **a,** RNA expression heatmap of intrinsic pathway pro-apoptotic genes (“intrinsic gene siganture”) from pre-treatment bone marrow samples. **b,** Integrated gene set score of intrinsic apoptotic signatures in patients with durable complete remission (CR) and non-responders (NR). **c,** Death receptor signature score adjusted for marrow blast counts prior to treatment in CRs and NRs. **d-e,** Pharmacokinetic analysis of tisagenlecleucel in patient peripheral blood over the first **c,** 90 and **d,** 1000 days after infusion, as measured by quantitative PCR of CAR transcripts grouped by marrow blast count-adjusted gene set score. **e,** Overall survival of all patients analyzed, grouped by adjusted death receptor signature score.

